# *De novo* genome and transcriptome analyses provide insights into the biology of the trematode human parasite *Fasciolopsis buski*

**DOI:** 10.1101/354456

**Authors:** Devendra K. Biswal, Tanmoy Roychowdhury, Priyatama Pandey, Veena Tandon

**Author notes:** Equal contributors (Devendra K. Biswal, Tanmoy Roychowdhury). Email addresses: DKB, TR, PP, VT.

## Abstract

Many trematode parasites cause infection in humans and are thought to be a major public health problem. Their ecological diversity in different regions provides challenging questions on evolution of these organisms. In this report, we perform transcriptome analysis of the giant intestinal fluke, *Fasciolopsis buski*, using next generation sequencing technology. Short read sequences derived from polyA containing RNA of this organism were assembled into 30677 unigenes that led to the annotation of 12380 genes. Annotation of the assembled transcripts enabled insight into processes and pathways in the intestinal fluke, such as RNAi pathway and energy metabolism. The expressed kinome of the organism was characterized by identifying all protein kinases. We have also carried out whole genome sequencing and used the sequences to confirm absence of some of the genes, not observed in transcriptome data, such as genes involved in fatty acid biosynthetic pathway. Transcriptome data also helped us to identify some of the expressed transposable elements. Though many Long Interspersed elements (LINEs) were identified, only two Short Interspersed Elements (SINEs) were visible. Overall transcriptome and draft genome analysis of *F. buski* helped us to characterize some its important biological characteristics and provided enormous resources for development of a suitable diagnostic system and anti-parasitic therapeutic molecules.

## Introduction

*Fasciolopsis buski* (giant intestinal fluke) is one of the largest digenean trematode flatworms infecting humans causing the disease fasciolopsiasis, with epidemiological records from Asia including the Indian subcontinent. Clinical manifestation of the disease is diarrhea, with presence of ulceration, intestinal wall abscess and hemorrhage, with the possibility of death if not treated in time [1]. Majority of the infected people (up to 60% in India and the mainland China) remain asymptomatic without manifestation of clinical invasive disease [1,2].

In India, these trematodes have been reported from different regions including the Northeast, associated with animal rearing (mainly pigs), or in underwater vegetables such as water chestnut. Although *F. buski* inhabits warm and moist regions and has been reported as single species in the genus, morphological variations among flukes from different geographical isolates have been observed suggesting genetic polymorphism or host-specific parasite adaptation [3,4].

Infective metacercarial cysts usually occur in 15-20 groups on the surface of aquatic vegetation [5,6] and once consumed, the juvenile adult stage of *F. buski* emerges and adheres to the small intestine of its host, attached until the host dies or is removed [1,2,7]. Unembryonated eggs are discharged into the intestine in the fecal material, with release of parasitic organisms (miracidia) two-week post infection. These hatch between 27–32°C, and invade snails (intermediate host) where developmental stages (sporocysts, rediae, and cercariae) are reported. The cercariae released and encysted as metacercariae (on aquatic plants), infect the mammalian host and develop into adult flukes completing its life cycle [8].

The current strategy for control of fasciolopsiasis, similar to that for other trematodes, such as *Gastrodiscoides hominis* is by blocking transmission among different hosts, i.e., human, animal reservoir and intermediate host (molluscan) [5]. Many drugs have been used to treat fasciolopsiasis. The drug of choice is praziquantel since it has high efficacy even in cases of severe fasciolopsiasis [9, 10, 11, 12, 13, 14]. More recently, the efficacy of triclabendazole, oxyclozanide and rafoxanide has been evaluated in pigs with favorable response to treatment [15]. Despite control programs, public health is a concern in endemic areas, and there is a need for new control measures with reports of sporadic re-emergence of infection (in Uttar Pradesh, India) from non-endemic regions [16, 17,18, 19].

Development of novel therapeutic agents targeting intestinal flukes is complex. The parasite biology occurs in multi-host life cycles and mechanisms for parasite host immune evasion strategies are not well understood. Genomics approaches are likely to be more successful in identifying new targets for intervention. Unfortunately, little genomic information is available for *F. buski* in the public domain other than a PCR-based molecular characterization using ITS 1 & 2 regions of ribosomal DNA (rDNA) genes [20,21]. Developments in next generation sequencing (NGS) technologies and computational analysis tools enable rapid data generation to decipher organismal biology [22, 23, 24, 25, 26]. NGS analysis to understand transcriptomes of related organisms, such as *Fasciola gigantica*, *Fasciola hepatica*, *Schistosoma mansoni*, *Schistosoma japonicum*, *Clonorchis sinensis*, *Opisthorchis viverrini*, *Fascioloides magna* and *Echinostoma caproni* reveal genes involved in adult parasite-host responses, such as antioxidants, heat shock proteins and cysteine proteases [27, 28, 29, 30, 31, 32]. Lack of molecular analysis of *F. buski* has hampered understanding of evolutionary placement of this organism among other *Fasciola*, as well as our understanding of mechanisms that regulate host-pathogen relationships. Previously we have reported the mitochondrial DNA (mt DNA) of the intestinal fluke for the first time that would help investigate Fasciolidae taxonomy and systematics with the aid of mtDNA NGS data [33]. Here, we report results of our effort to characterize the transcriptome of the adult stage of *F. buski*. NGS technology was used to get RNA-seq and bioinformatics analysis was carried out on the sequence output. In a few cases we have used the *F. buski* draft genome sequence to confirm our observations.

## Materials and methods

### Source of parasite material

*F. buski* adult specimens were obtained from the intestine of freshly slaughtered pig (*Sus scrofa* domestica) by screening for naturally infected pigs among the animals routinely slaughtered for meat purpose at the local abattoirs. The worms reported in this study represent geographical isolates from the Shillong area (25.57°N, 91.88°E) in the state of Meghalaya, Northeast India. Eggs were obtained by squeezing mature adult flukes between two glass slides. Adult flukes collected from different pig hosts were processed singly for the purpose of DNA extraction and eggs were recovered from each of these specimens separately.

All sample were obtained from animals used for local meat consumption and no live animals were handled or sacrificed for this study. Hence there was no need to follow relevant guidelines as laid down for handing live vertebrates.

### RNA and DNA extraction, Illumina sequencing

Briefly, the adult flukes were digested overnight in extraction buffer [containing 1% sodium dodecyl sulfate (SDS), 25 mg Proteinase K] at 37°C, prior to DNA recovery. Genomic DNA was then extracted from lysed individual worms by ethanol precipitation technique or by FTA cards using Whatman’s FTA Purification Reagent [34]. Quality and quantity of the DNA was assessed by 0.8% agarose gel electrophoresis (Sigma-Aldrich, USA), spectrophotometer (Nanodrop 1000, Thermofischer USA) and flurometer (Qubit, Thermofischer, USA). Whole genome shot-gun libraries for Illumina sequencing were generated and the genomic DNA was sheared (Covaris, USA), end-repaired, adenylated phosphorylate ligated to Illumina adapters (TruSeq DNA, Illumina, USA), to generate short (300-350 bp) fragment and long (500 – 550) bp insert libraries. Both libraries were amplified using 10 cycles of PCR KapaHiFiHotstart PCR ready mix (Kapa Biosystems Inc., USA) and adapter removal by Solid Phase Reversible Immobilization (SPRI) beads (Ampure XP beads, Beckmann-coulter, USA). Prior to sequencing, fragment analysis was performed (High Sensitivity Bioanalyzer Chip, Agilent, USA), quantified by qPCR. 36cycle sequencing was performed on Illumina Genome analyzer II (SBS kit v5, Illumina, USA) and Illumina HiSeq1000 (Illumina, USA) to generate 16 million 100 paired-end reads from the shot-gun library.

Total RNA was collected from a single frozen adult individual fluke (~100mg tissue) using TRI reagent^®^ (Sigma, USA). The integrity of total RNA was verified using Bioanalyzer Agilent 2100 with RNA integrity number 9. Transcriptome libraries for sequencing was constructed from poly A RNA isolated from 1ug total RNA (OligoTex mRNA minikit, Qiagen Gmbh, Germany), that was fragmented, reverse transcribed (Superscript II Reverse transcriptase, Invitrogen, USA) with random hexamers as described in (TruSeq RNA Sample Preparation Kit, Illumina, USA). The cDNA was end-repaired, adenlylated and cleaned up by SPRI beads (Ampure XP, Beckman Coulter, USA) prior to ligation with Illumina single index sequencing adapters (TruSeq RNA, Illumina, USA). The library was amplified using 11 cycles of PCR and quantified using Nanodrop (Agilent). The prepared library was validated for quality by running an aliquot on High Sensitivity Bioanalyzer Chip (Agilent) that displayed a confident bioanalyzer profile for transcriptome sequencing.

### De novo assembly, annotation and characterization of transcriptome

Illumina-generated paired end reads were filtered for low quality scores (PHRED score < 30). From 32010511 (32.01 millions) high quality raw reads (>70% of bases in a read with >20 phred score), 47957 contiguous sequences were assembled. Reads containing ambiguous characters were also removed from the dataset. The high-quality reads were then assembled using Trinity software [35] designed for transcriptome assembly. Default parameters were used for this purpose. While Trinity can identify different isoforms, our initial target was to identify the unigene transcripts. Thus, transcripts were subjected to clustering using CD-HIT-EST [36] at 90% similarity. To assess the quality of assembly, raw reads were mapped back to unigenes using Bowtie [37]. Reads aligning in multiple locations were randomly allocated to one of the unigenes. Different properties, such as average read depth, coverage, and percent uniquely mapped reads were calculated. The numbers of raw reads were normalized for length and RPKM (reads per kilobase of exon per million of mapped reads) values were calculated in order to quantify relative abundance of each unigene for estimating the levels of expression.

All unigenes were annotated using both sequence and domain-based comparisons that involved sequence similarity analysis with non-redundant (NR) protein database of NCBI (http://www.ncbi.nlm.nih.gov/), Uniprot (Swissprot + trEMBL) [38] and Nembase4 EST database [39] using BLASTx and tBLASTn [40], Significant threshold of E-value ≤ 10^−5^ was used as a cutoff. Since we were not able to annotate large numbers of unigenes using sequence similarity, conserved domains were further identified in InterPro database [41] (HMMpfam, HMMsmart, HMMpanther, FPrintScan, BlastProDom) and Clusters of Orthologous Groups (COG) database [42] using InterProScan and BLASTx respectively. Functional categorization was assigned by finding Gene Ontology terms of best BLASTx hits against NR database using Blast2GO software [43] (http://www.blast2go.com/b2ghome). GO assignments were represented in a plot generated by WEGO [44]. Comparative analysis of the transcriptome was performed against other trematodes. Transcriptome datasets of trematode parasites, viz. *Fasciola hepatica*, *Fasciola gigantica*, *Clonorchis sinensis*, *Opisthorchis viverrini*, *Schistosoma mansoni* and *Schistosoma japonicum* were downloaded from NCBI Short Read Archive (SRA) database (http://www.ncbi.nlm.nih.gov/sra/). Unigenes were mapped to different KEGG pathways using KAAS [45]. All unigenes were used against the KEGG database by a Bi-directional best-hit method suitable for whole genome or transcriptome data. KEGG orthologs were displayed in iPath2 [46]. ORFs were predicted using ORFpredictor [47] (http://proteomics.ysu.edu/tools/OrfPredictor.html). Signal peptides were identified using SignalP [48] (http://www.cbs.dtu.dk/services/SignalP/) and transmembrane domains were identified using TMHMM [49]. Kinases and peptidases were identified by comparison against EMBL kinase database (http://www.sarfari.org/kinasesarfari) and MEROPS database [50] consecutively. Orthologs of RNAi pathway proteins of *Caenorhabditis elegans* in *F.buski* were identified by a reciprocal BLAST hit strategy. Raw reads generated from *F. buski* transcriptome were submitted in NCBI SRA (https://trace.ncbi.nlm.nih.gov/Traces/sra/?run=SRR941773) under the accession number SRX326786.

### *De novo* assembly of draft genome and identification of transposable elements

Short read sequences from the *F. buski* genome were assembled using a number of assembly tools, such as SOAP denovo [51], Velvet [52] and Abyss [53]. These algorithms use de Bruijn graph for genomic sequence assembly. Default parameters were used to assemble 42425988 paired end reads of different insert size libraries. Each assembly program was used with a range of K-mers. Statistics of best assemblies in respect of N50 values by three programs are given in S1 Table. These assembly results suggest Trinity assembler generated higher N50 comparatively for this dataset. Further analysis of *F. buski* genome was performed on this reference draft genome assembly. RepeatMasker [54] was used to identify repetitive and transposable elements with default parameters. It makes use of RepBase libraries [version 20120418] (www.girinst.org) that are used as a reference for the identification of transposable and repetitive elements in a query sequence. Transcripts obtained from the transcriptome were matched to the draft assembly. The raw reads generated from *F. buski* genome were submitted in NCBI SRA (www.ncbi.nlm.nih.gov/sra/) under the accession numbers SRX3087506, SRX327895 (genome data) and SRX326786 (transcriptome data).

## Results

### *De novo* assembly and annotation of transcriptome

The whole genome and transcriptome data are sequenced from the adult specimens of *F. buski* that were collected from pigs with a naturally acquired infection at an abattoir in Shillong, Meghalaya. Detailed statistics of initial output generated by Trinity is shown in S1 Table. Initial output was then subjected to CD-HIT-EST to identify clusters and the results showed 43111 clusters with 3298 singletons representing 46409 unigene transcripts. After assessment of the assembly (S2 Table), 30677 high-quality unigenes were used for subsequent analysis. We were able to predict ORFs for 30628 unigenes with the mean length of ORF being 185 amino acids. The methionine start codon (ATG) was present in 23759 (77.4%) ORFs.

*F. buski* unigenes matched with 12279 sequences of NCBI non-redundant (NR) database with an E-value less than 1E-05 on BLAST analysis (S3 Table). Best hit for each unigene was ranked according to E-values. We report many significant BLAST hits to *C. sinensis* (55%), *S. mansoni* (21%) and *S. japonicum* (14%), with less similarity to *F. hepatica* or *F. gigantica*. This may be due to the availability and skew of sequence databases for the two *Fasciola* species. Although, an identity cutoff of E-value ≤ 1E^−05^ was applied for identification of remote homologs, 68% of the unigenes were matched with similarity of E-values less than 1E^−30^. Annotation of the unigenes depended on the length of the unigenes, and shorter unigenes (13% of smaller than 500 bp) were not matched, while the longer unigenes (91% of unigenes above 2000 bp) were annotated. The high percentage of annotation suggested an accurate assembly, and presence of functionally informative datasets for *F. buski*.

Unigenes were also annotated by sequence comparison against Uniprot/Swissprot and Uniprot/trEMBL database. We found 8089 unigenes that showed significant similarity with at least one Swissprot entry and 12231 with at least one trEMBL entry, totaling 12242 unique unigenes. Analysis using NEMBASE4 database helped us to find putative function of 6776 unigenes. Overall 6752 unigenes displayed significant sequence matches against all three databases used for sequence-based annotation.

Identification of conserved domains and motifs sometimes helps in functional classification of genes. Interpro scan helped us to identify 3309 unique domains/families in 6588 unigenes. Out of these, 1775 (53.6%) domains/families were associated with only one unigene. Some of the best hits are given in S4 Table. Among these, protein kinases and ankyrin repeat domains were found to be more prevalent. Protein kinases play a significant role in signaling; Ankyrin repeat containing proteins are involved in a number of functions, such as integrity of plasma membrane, and WD40 repeat is associated with signal transduction and transcription regulation.

Unigenes were also classified using Clusters of Orthologous (COG) database at http://www.ncbi.nlm.nih.gov/COG/. Each entry in COG is supported by the presence of the sequence in at least three distinct lineages, thereby representing an ancient conserved domain. A total of 4665 (15.2%) unigenes were assigned with different functional terms (S5 Table). Though most of the unigenes displayed more than one hit, a majority of them were from the same COG functional class. Best hits were further processed for functional classification. The results are shown in Fig 1. Large numbers of unigenes mapped to “Post translational modification, protein turn over, chaperons” (354; 7.5%), “Translation, ribosomal structure and biogenesis” (289; 6.1%), “Replication, recombination and repair” (216; 4.6%) and “Signal transduction mechanisms” (195; 4.1%) in addition to “general function prediction” (938; 20.1%). In contrast, there were very few genes that mapped to functional classes “cell motility” (7) and “nuclear structure” (2).

**Fig 1.**
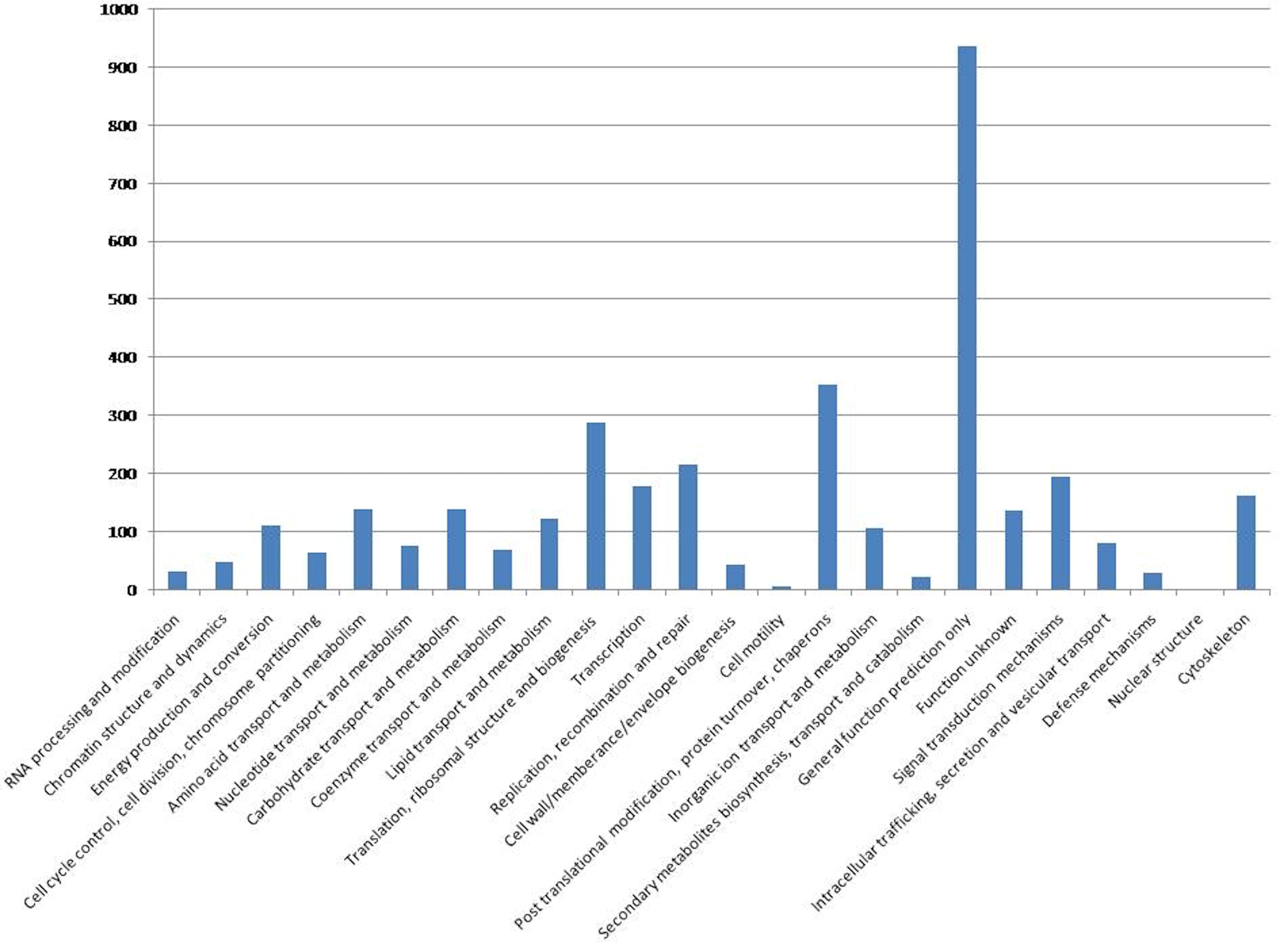
COG classification of *F. buski* transcriptome. The X-axis and Y-axis represent different COG categories and number of unigenes associated with each COG category respectively.

Since secretory proteins, such as those that are classified as excretory and secretory proteins (ES), are involved in host-parasite interactions, we identified proteins that belong to this category. Our analysis showed that 1501 unigenes contained signal peptides and there were 1406 ES proteins. Their identification was done by the presence of a signal peptide domain and the absence of any trans-membrane domain.

Sequences that matched significantly with NCBI nr database were further processed using BLAST2GO for GO assignment. This analysis was done to further classify expressed genes into different functional classes. A total 36433 GO terms were assigned to 6658 unigenes. The results of this analysis are depicted in Fig 2. Majority of the GO terms were from “biological processes” (2627; 56.2%), while others represented “molecular function” (1445; 30.9%) and “cellular component” (602; 12.8%). The major sub-categories were found to be mostly those associated with cell structure and function and metabolic processes, such as “cell” (GO: 005623), “cell part” (GO: 0044464), “macro-molecular complex” (GO: 0032991) and “organelle” (GO: 0043226), “catalytic function” (GO: 0003824) and “metabolic process” (GO: 0008152). Considering all levels of GO assignment, a significant number of sequences were found to be associated with “Regulation of transcription, DNA-dependent” (GO: 0006355; 344), “integral to membrane” (GO: 0016021; 925) and “ATP binding” (GO: 0005524; 1086).

**Fig 2.**
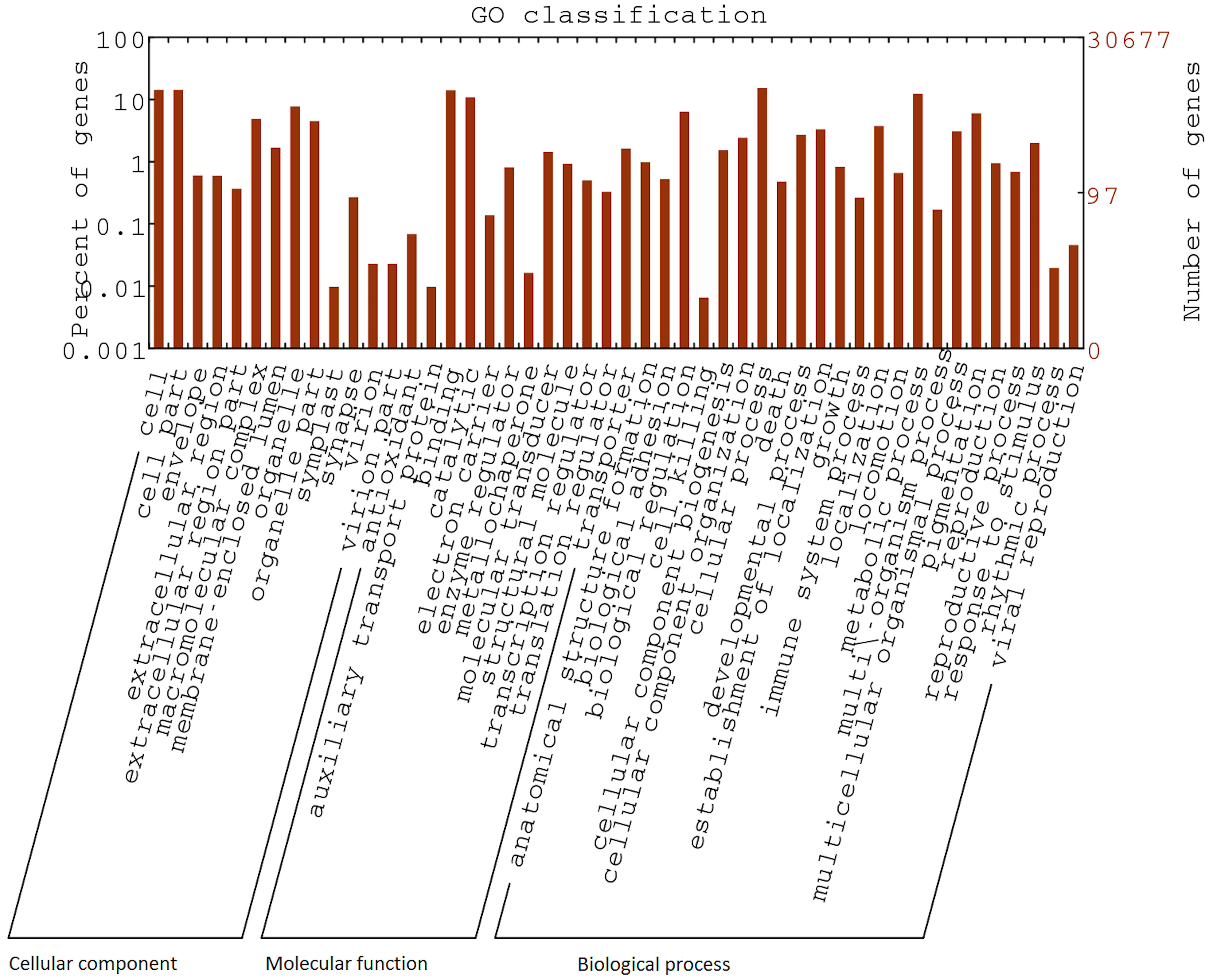
Gene ontology classification of *F. buski* transcriptome. GO terms assigned to Unigenes were classified into three major functional classes; Cellular component, Biological process and Molecular function.

A Venn diagram summarizing annotation analysis using five databases is given in Fig 3. It was possible for us to annotate 12380 unigenes using both sequence as well as domain-based comparisons. The mean length of annotated unigenes was found to be 1727, which is higher than the mean length (1080) of all the unigenes thus, showing the importance of assembly quality for annotation (S1 Fig). It is expected that unannotated unigenes may participate in unique biological functions specific to this organism.

**Fig 3.**
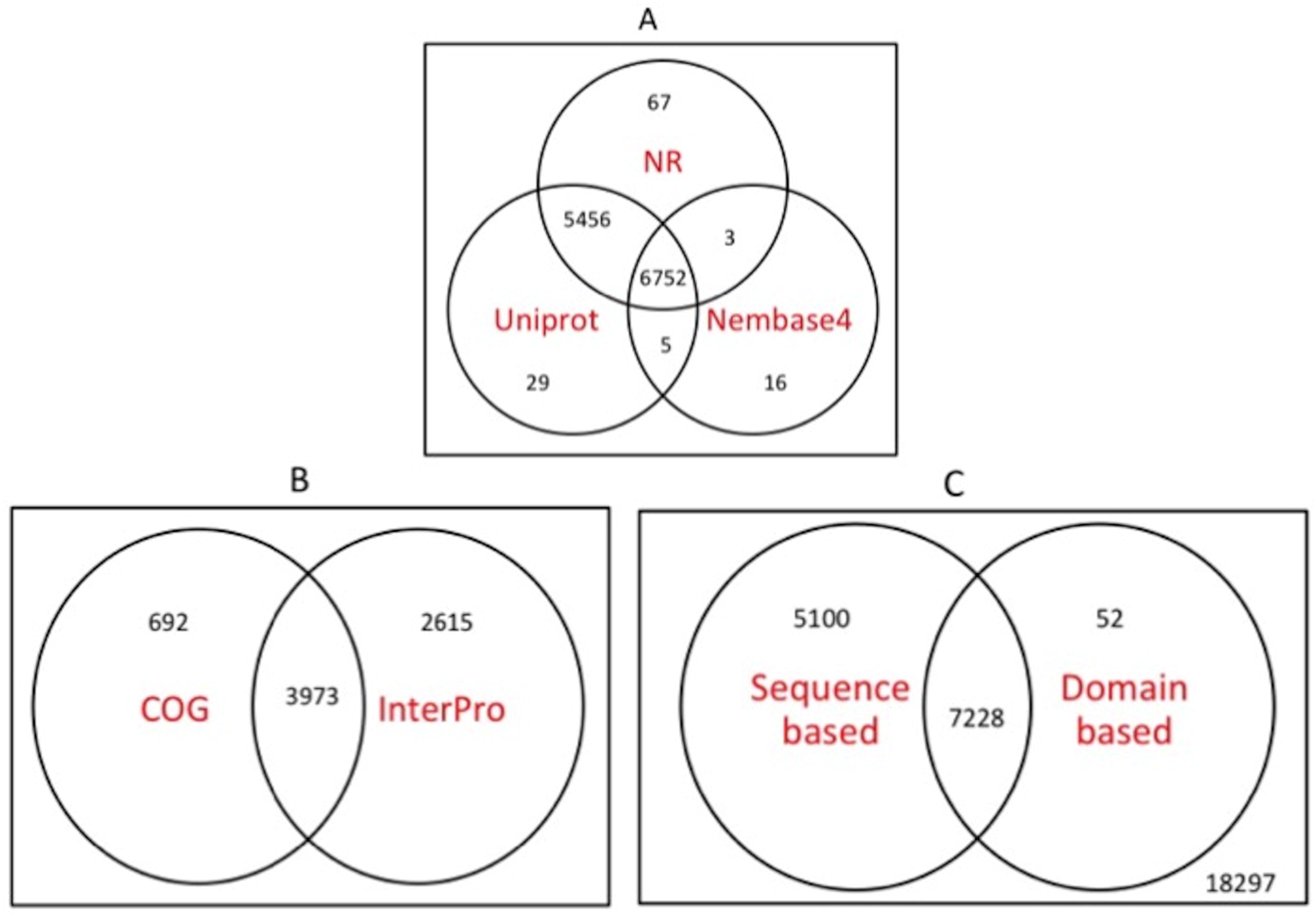
Venn diagram showing a number of annotated unigenes using sequence based and domain based comparisons. A) Unigenes were compared with NR, Uniprot and Nembase4 for sequence-based annotation. Circle intersections represent number of unigenes found in more than one databases. B) Unigenes were compared against COG and INTERPRO databases for domain-based annotation. C) Two circles A and B represent number of sequences annotated using sequence based and domain based comparisons respectively. The numbers outside circles represent number of sequences remained unannotated after employing all these comparisons.

### Comparison with other trematodes

A comparative sequence analysis was performed with RNA-seq datasets of different trematode families in order to understand functional divergence. The results are summarized in Table 1 and in Venn diagram (Fig 4). The analysis revealed that 54% of the unigenes displayed sequence similarity with one of the trematodes and 19% with all trematodes analyzed at an E-value threshold of 1E^−05^ When sequence similarity values at different E-value cut-offs were compared transcript sequences from *F. gigantica* appeared to be evolutionarily the closest as it displayed highest number of matches at a given E-value among trematodes studied. Further, to reconstruct evolutionary relationships of *F. buski* with other trematodes, we computed a phylogenetic tree (Fig 5) concatenating all the annotated 12 protein-coding genes (PCGs) from the *F. buski* mitochondrial DNA (mtDNA). The mtDNA for the intestinal fluke was recovered from the *F. buski* genome data and is available in NCBI SRA bearing accession no. SRX316736. Previous studies confirm that alignments with > 10,000 nucleotides from organelle genomes have ample phylogenetic signals for evolutionary reconstruction of tapeworm phylogeny [55]. The phylogenetic tree is well supported by very high posterior probabilities as the taxa grouped well into distinct clades representing the different worm Families such as Opisthorchiidae, Paragonimidae, Paramphistomidae and Fasciolidae (Trematoda); Ascarididae (Nematoda) and Taeniidae (Cestoda). *F. buski* claded well with *F. hepatica* and *F. gigantica* with strong bootstrap support.

**Table 1.**
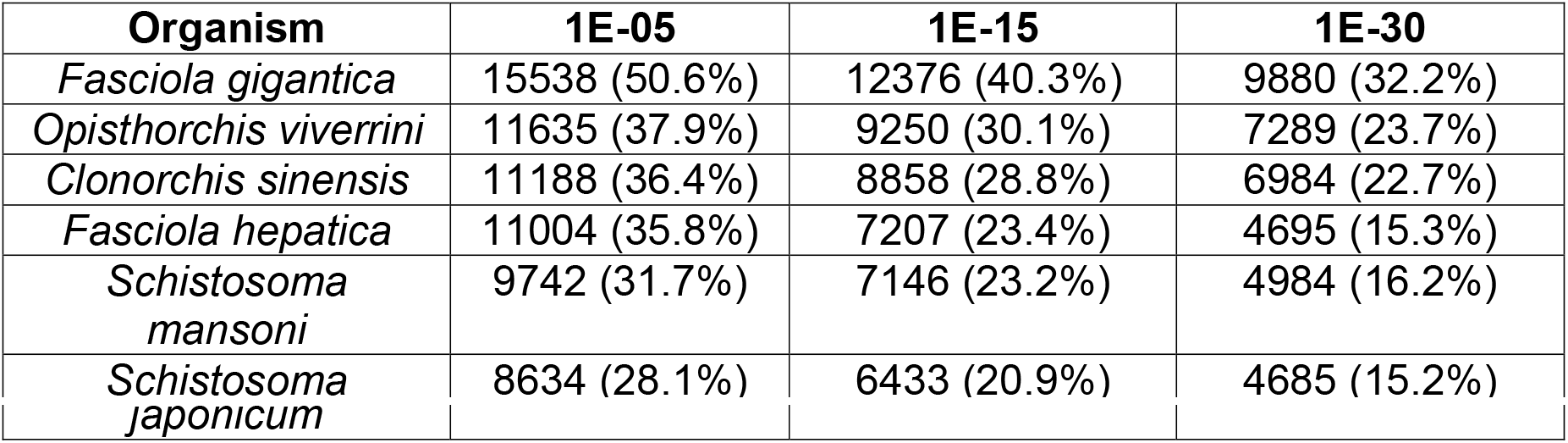
Comparative sequence analysis of *F. buski* transcriptome with different trematodes.

**Fig 4.**
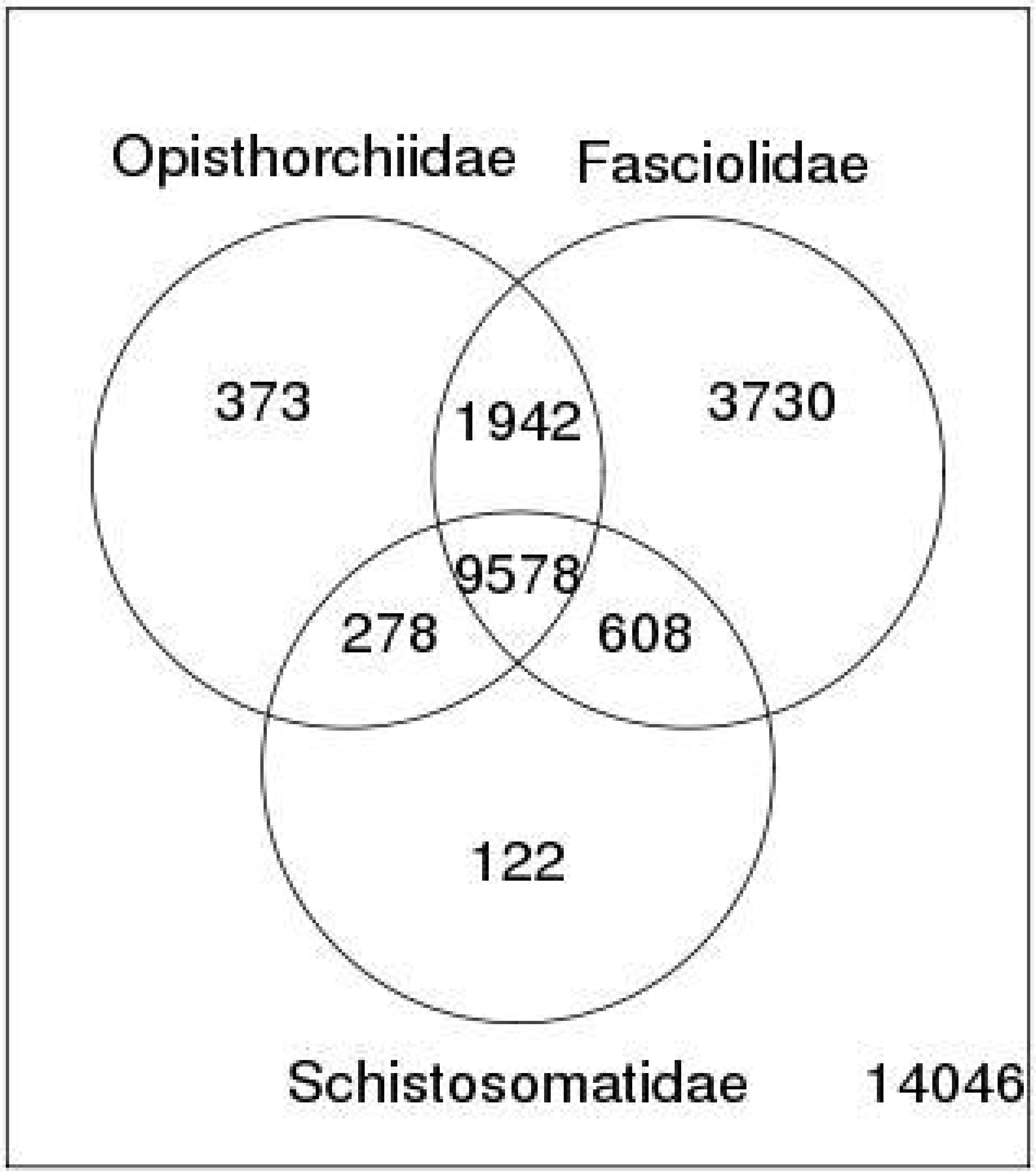
Venn diagram showing number of homologs found in *F. buski* transcriptome against transcriptome/EST datasets of major trematode families of Opisthorchiidae (*Opisthorchis viverrini* and *Clonorchis sinensis*), Fasciolidae (*Fasciola hepatica* and *Fasciola gigantica*) and Schistosomatidae (*Schistosoma mansoni* and *Schistosoma japonicum*).

**Fig 5.**
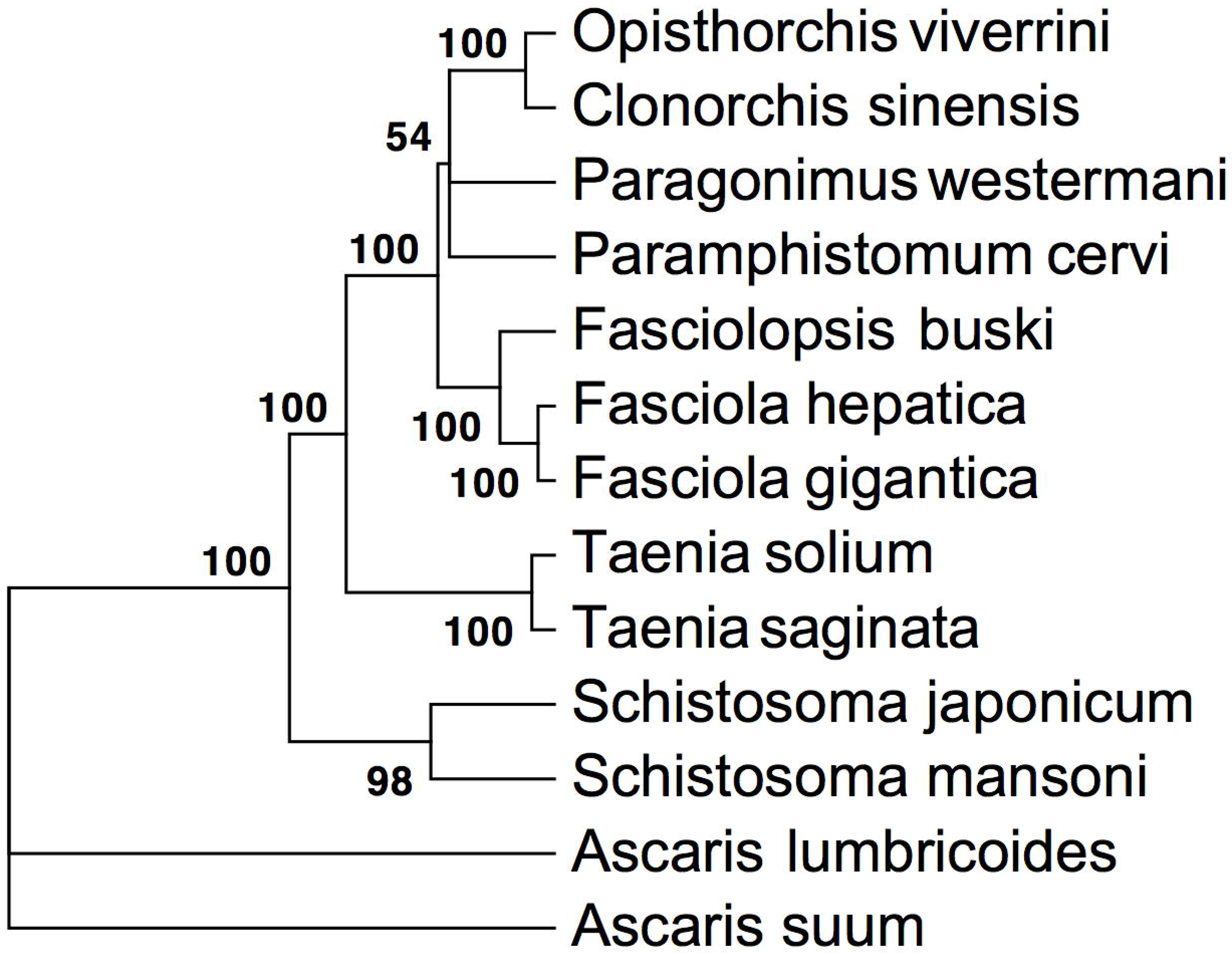
Bayesian phylogenetic relationship among the representative helminth species based on 12 PCGs from their mitochondrial DNA. Phylogenetic analyses of concatenated nucleotide sequence datasets for all 12 PCGs were performed using four MCMC chains in bayesian analysis run for 1,000,000 generations, sampled every 1,000 generations. Bayesian posterior probability (BPP) values were determined after discarding first 25% of trees as burn-in. Posterior support values appear at nodes. Species representing Nematoda (Ascaridida) were taken as outgroup.

Out of the homologs of *F. buski* unigenes found in trematodes but not in model eukaryotes, 3552 could be annotated. These genes are likely to be trematode specific and can be a useful resource in development of new diagnostic and therapeutic tools. Our results suggest relatively high level of sequence divergence among coding regions (ortholog divergence) of different trematodes, probably a result of adaptation to different ecological conditions.

### Pathway analysis

We used KEGG automatic annotation server (KAAS) for identification of pathways associated with unigenes. About 3026 sequences were annotated with 2527 KO terms. In total, 287 different pathways, classified into six major groups, were identified (Table 2). Some of the genes could not be tracked in the transcriptome. This is presumably due to lack of expression in the adult flukes, but could also represent transcripts that are not polyadenylated and thus were not included in our transcript pools. In either case, a high quality genome sequence data with better depth coverage would be required to confirm the existence or absence of the genes. Major pathways deciphered are those involved in metabolism (638 unigenes, 520 KO terms), such as carbohydrates, amino acids, energy metabolism and genetic information processing (790 unigenes, 690 KO terms) transcription, translation, replication and repair, and those involved in human diseases (530 unigenes, 440 KO terms) cancer and neurodegenerative disorders. Overall, most highly represented KEGG terms by virtue of number of unique KO identifiers are Spliceosome (33), RNA transport (92) and protein processing in endoplasmic reticulum (83). Major components of metabolic pathways found in *F. buski* transcriptome are shown in S2 Fig. We could not detect most of the components of fatty acid biosynthesis pathway except acetyl-CoA/propionyl-CoA carboxylase [EC: 6.4.1.2; EC: 6.4.1.3] and 3- oxoacyl-[acyl-carrier-protein] synthase II [EC: 2.3.1.179]. We could not also detect these genes in the current draft genome of *F. buski*. Probably this can be better resolved with an improved genome assembly with higher depth coverage. While EC: 2.3.1.179 was identified in *F. hepatica*, *C. sinensis*, *O. viverrini*, *S. mansoni* transcriptomes [27, 28]; it was reported to be missing in *F. gigantica* [29]. It is not surprising as these parasites acquire fatty acids from the host [56]. Pathways encoding enzymes of different amino-acid-biosynthesis pathways were represented only by one enzyme for valine, leucine and isoleucine biosynthesis [map00290], and under-represented for pathways, such as lysine biosynthesis [map00300] where no match was detected. In contrast, genes encoding enzymes in fatty acid metabolism [map00071] and amino acid metabolism [map00280; map00310; map00330] are well represented in the *F. buski* transcriptome. The skewed representation of enzymes in the pathway analysis provided clues on the gene regulation and parasite biology pointing towards catabolic process, with a likely dependence in this stage of its life cycle to their host for nutrition.

**Table 2.**
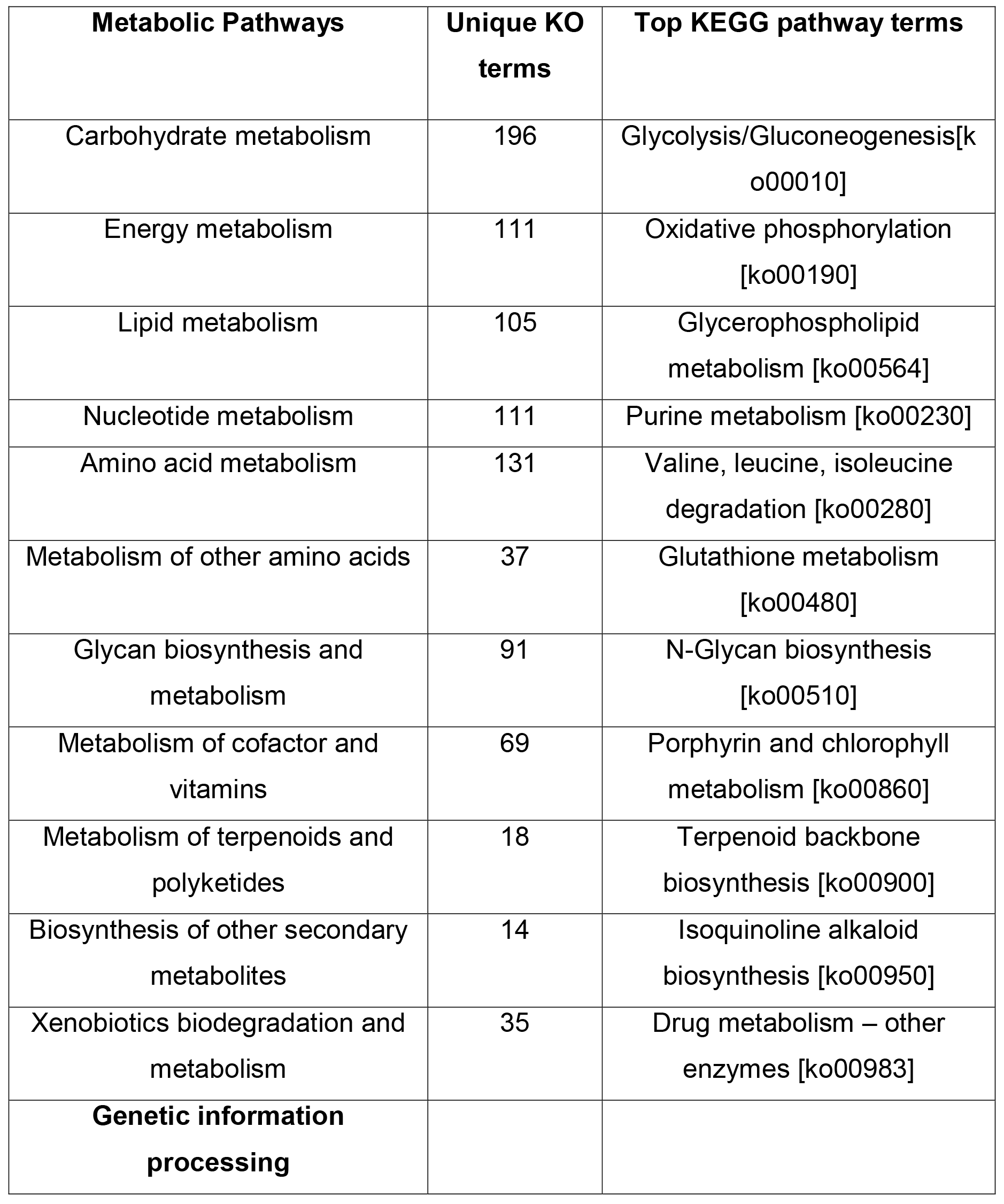
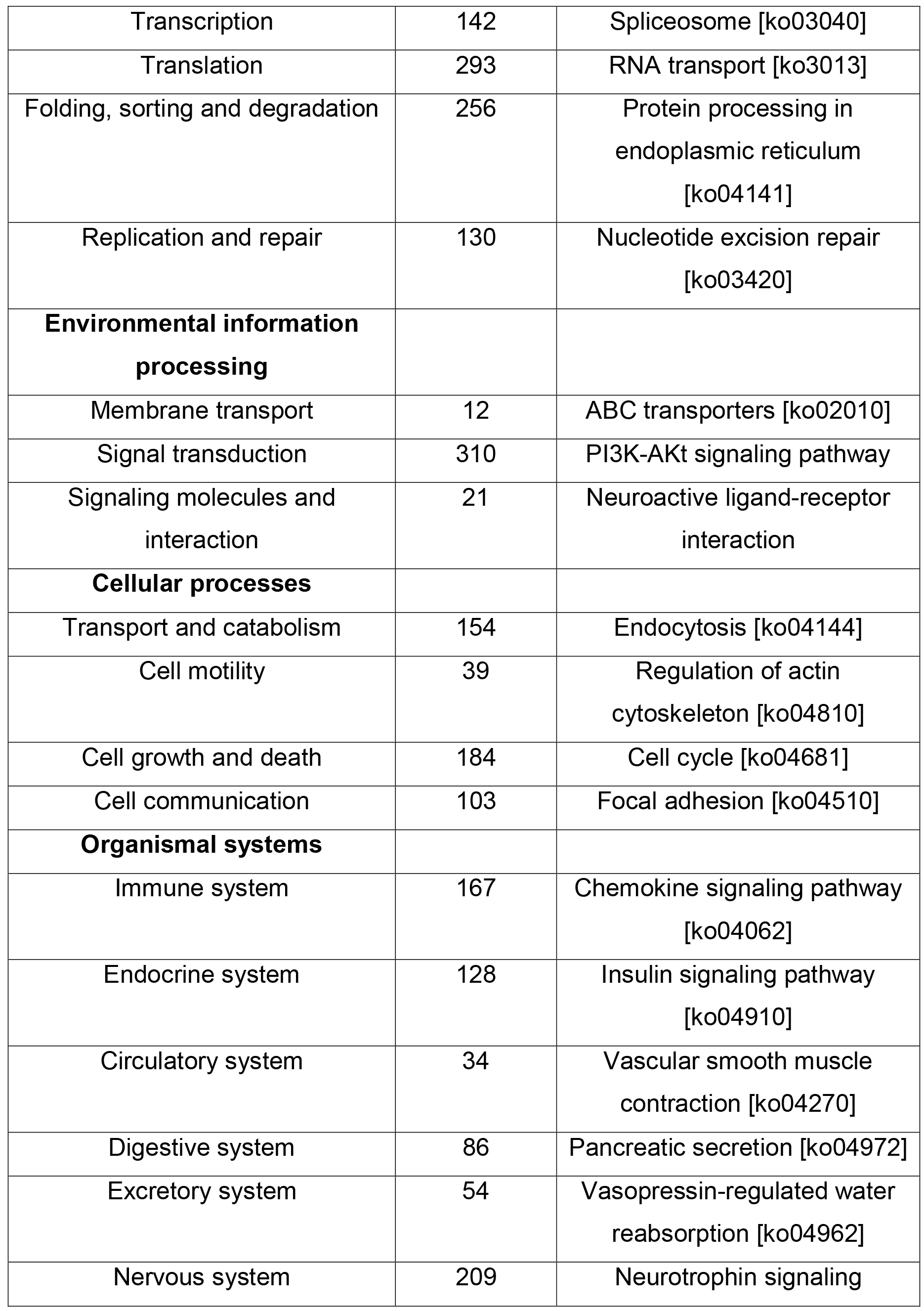
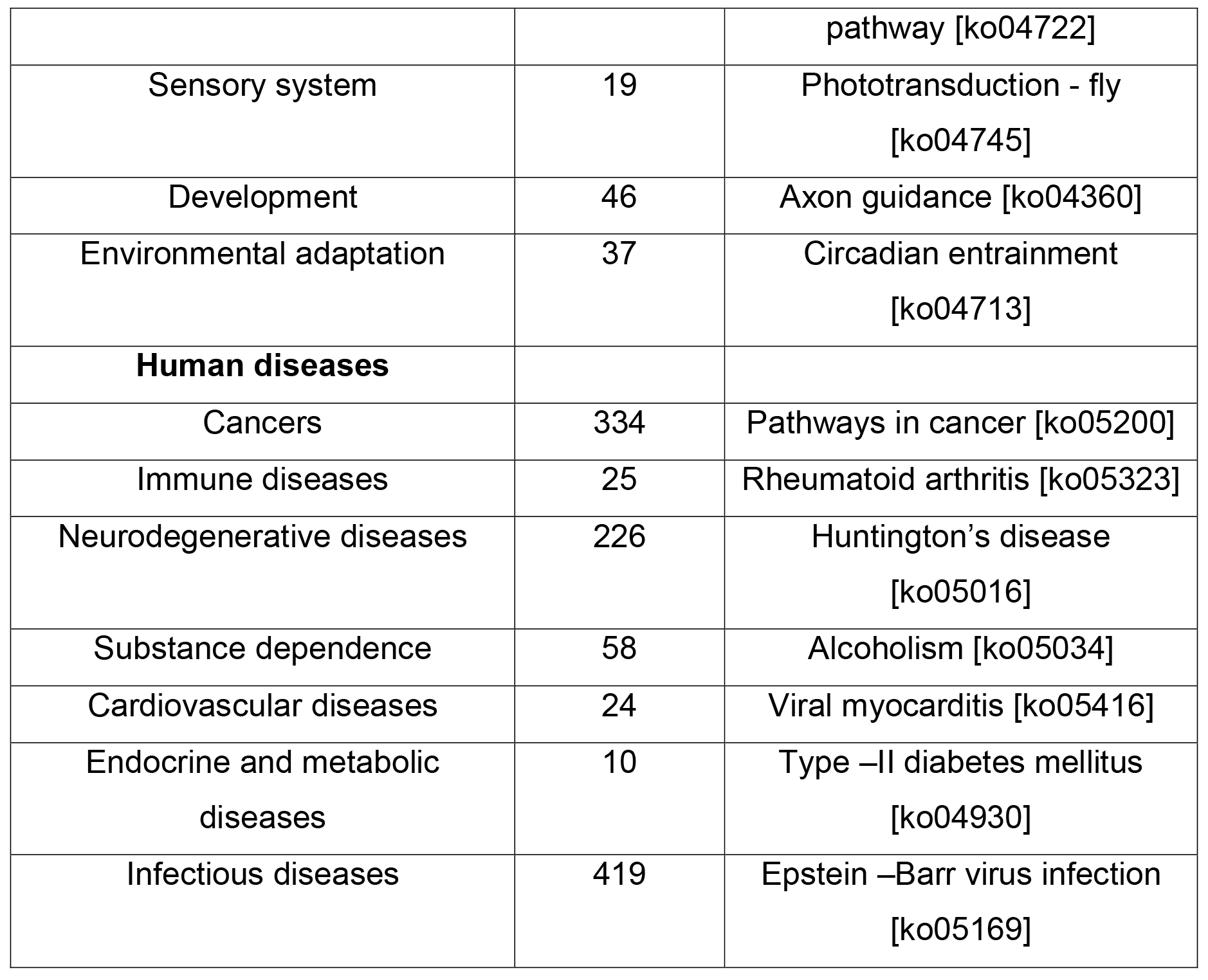
KEGG pathways identified from *F. buski* transcriptome using KAAS.

### Highly expressed genes

Expression status of transcripts can provide clues on their likely biological relevance [57]. To determine the relative expression levels RPKM values of each unigene was evaluated (S6 Table). RNA-seq can provide sensitive estimates of absolute gene expression variation [57]. We noted expression variation of unigenes upto 4th order of magnitude. However, only a few unigenes (5) displayed RPKM value >10,000 and 113 genes exceeding 1000. Among top 100 highly expressed genes, functions linked to translation (ribosomal genes-40S, 60S ribosomal proteins, Elongation Factor EF1α, Translation Initiation Factor TIF1), protein stress and folding (heat shock proteins 20 and 90 and 10 kDa chaperonins) constituted the majority. As expected, unigenes encoding cytoskeletal proteins were expressed at relatively high levels. Since proteases play an important role in parasite host-pathogen interaction, it was not surprising to see highly expressed Cathepsin L and protease inhibitors as part of the transcriptome. These observations are similar to that of *S. mansoni*, *F. gigantica*, *C. sinensis and O. viverrini* [27, 29, 58]. In these trematodes stress response genes, genes associated with ribosomes and translation, actin myosin complex and proteolytic enzymes appear to be highly expressed. The results suggest that the adult parasite is active in protein synthesis and is utilizing nutrients from the host to function at a high metabolic load for reproduction and egg development.

### Kinome

Eukaryotic or conventional protein kinases (ePKs) play important regulatory roles in diverse cellular processes, such as metabolism, transcription, cell cycle progression, apoptosis, and neuronal development [59]. These are classified into eight groups, based on sequence similarity of their catalytic domains, the presence of accessory domains, and their modes of regulation [60, 61, 62]. In addition to eight ePKs, a ninth group categorized as ‘Other’ and consisting of a mixed collection of kinases that cannot be classified easily into any of the groups is defined in the EMBL kinase database [41]. There are extensive studies on kinome of the model organism *C. elegans*. Kinases from this free-living nematode are deeply conserved in evolution, and the worm shares family homologs as high as 80% with the human kinome. Out of a total of 438 worm kinases nearly half are members of worm-specific or worm-expanded families. Studies point to recent evolution of such homologous genes in *C. briggsae* involved in spermatogenesis, chemosensation, Wnt signaling and FGF receptor-like kinases [63]. Our analysis of the *F. buski* transcriptome suggested 190 ePKs (250 unigenes) belonging to all the nine groups of protein kinases. The results are shown in Fig 6 and S7 Table. The presence of multiple protein kinase families, actively expressed in the adult stage suggests that *F. buski* encodes an extensive signaling network and majority of these eukaryotic signal transduction pathways are conserved. In *F. gigantica*, 308 sequences of protein kinases belonging to eight ePK classes were found in the transcriptome dataset [29], in contrast the free-living nematode *C. elegans* encoded nearly 400 pathways [64]. We ascribe this to the parasite biology or possible stage specific expression that could not be determined from the transcriptome dataset of single stages.

**Fig 6.**
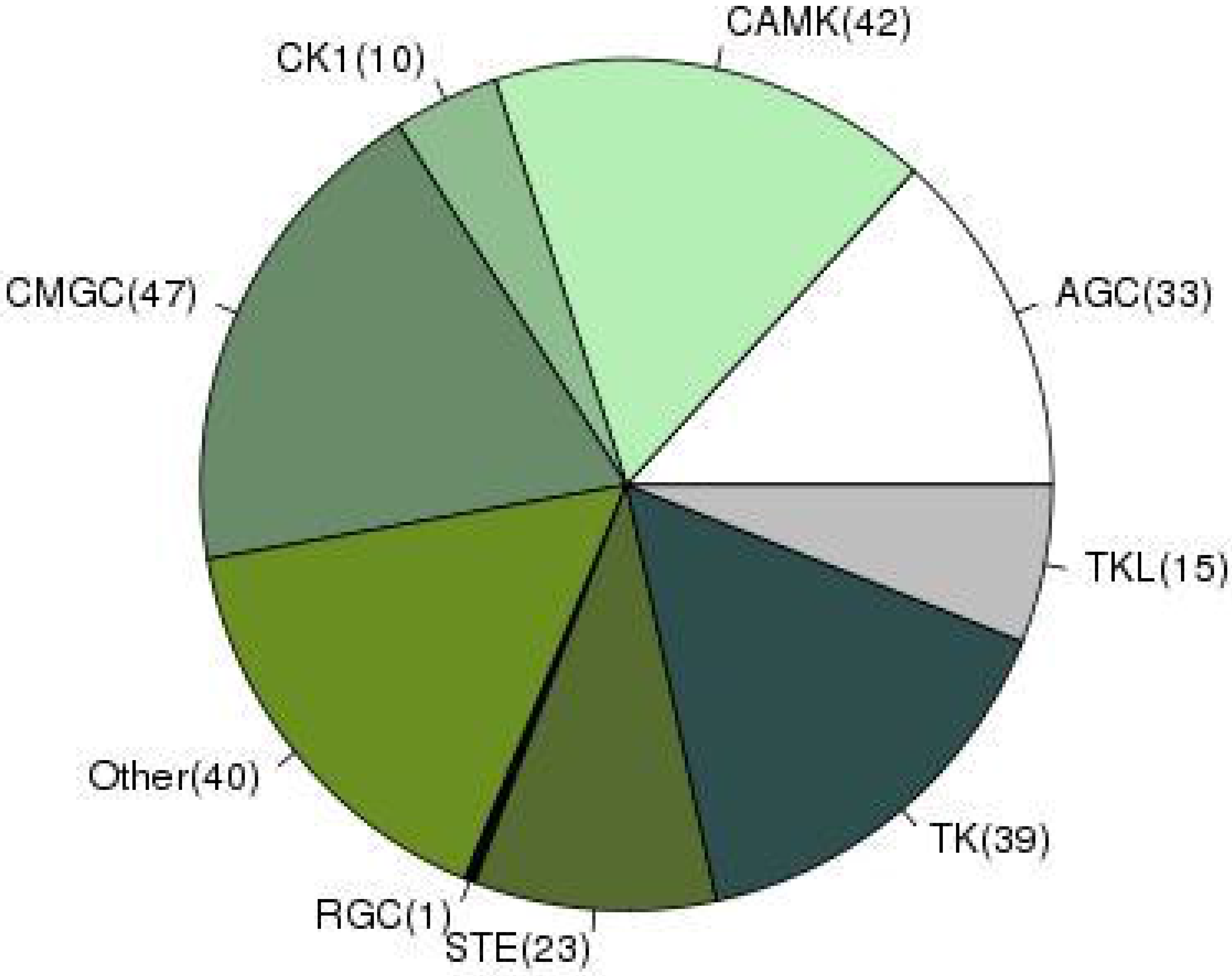
A pie chart displaying a number of significant matches found against nine different protein kinase classes (according to EMBL kinase database) in *F. buski* transcriptome. The protein kinase groups are represented by: (i) CMGC-cyclin-dependent, mitogen activated, glycogen synthase and CDK-like serine/threonine kinases; (ii) CAMK – Calcium/Calmodulin-dependent serine/threonine kinases; (iii) TK – Tyrosine kinases; (iv) TKL – Tyrosine kinase-like; (v) AGC – cAMP-dependent, cGMP-dependent and protein kinase C serine/threonine kinases; (vi) STE – serine/threonine protein kinases associated with MAP kinase cascade; (vii) CK1 – Casein kinases and close relatives; (viii) RGC – Receptor guanylate cyclase kinases: represented by a single protein, receptor guanylate cyclase kinase and (ix) other unclassified kinases.

### Protease and protease inhibitors

*F. buski* transcriptome suggests that a total of 478 proteases (MEROPS terms), corresponding to 6 catalytic types, and 138 protease inhibitors (138 MEROPS terms) as defined in the MEROPS database are expressed in this organism [50]. The results are shown in Figure 7. Some of the highly expressed proteases are prolyl oligopeptidase family **of** serine proteases (206 MEROPS terms; 16 families), lysostaphin subfamily **of** metalloproteases (126 MEROPS terms; 21 families) and ubiquitin-specific peptidases **of the** cysteine proteases family (81 MEROPS terms; 22 families) in addition to a significant number of protease inhibitors (138 MEROPS terms; 14 families) (Fig 7 and S8 Table)

**Figure 7:**
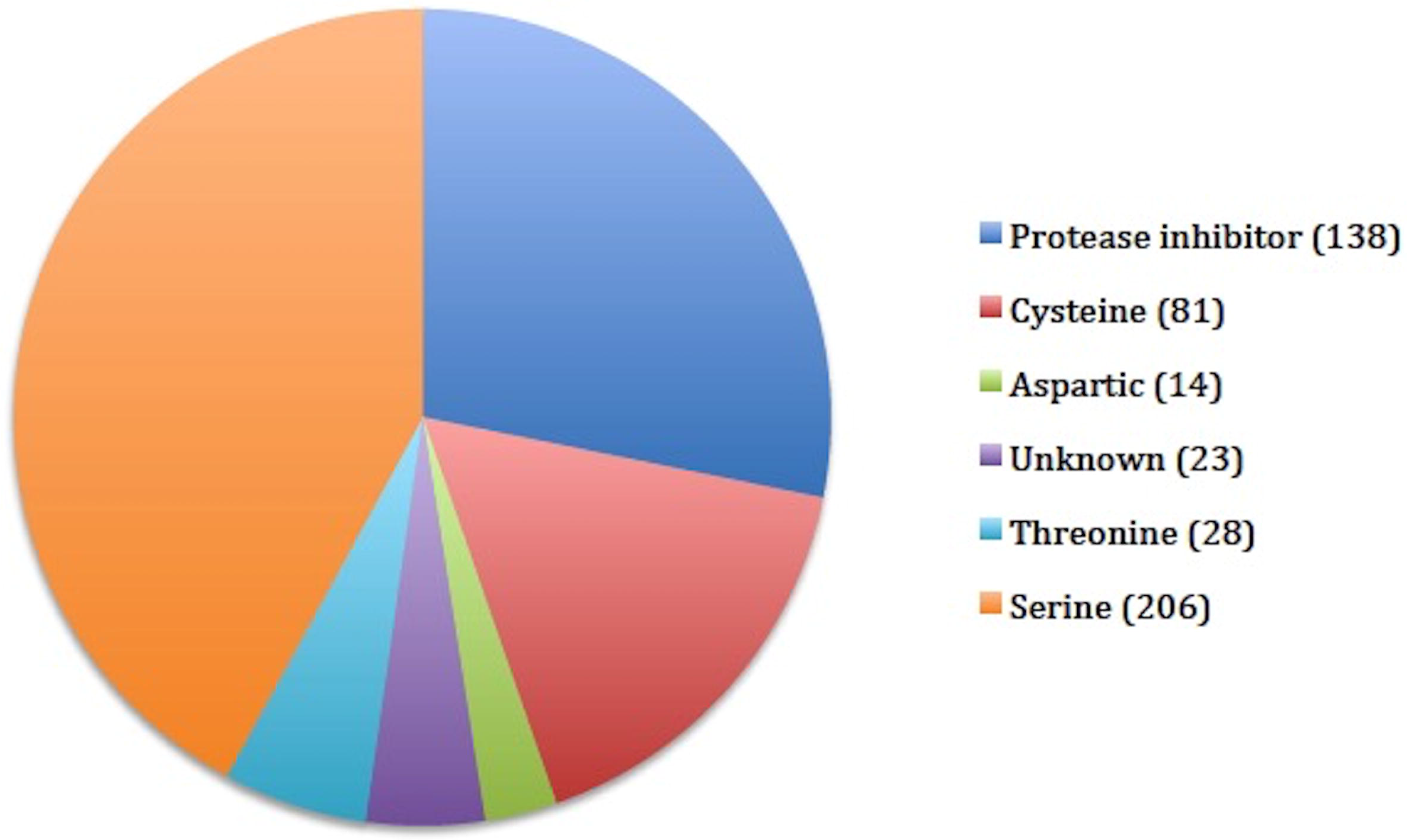
Protease and protease inhibitors (MEROPS terms) found in *F. buski* transcriptome.

### RNAi pathway genes

RNA interference (RNAi), a gene silencing process generally triggered by double-stranded RNA (dsRNA) delivers gene-specific dsRNA to a competent cell, thereby engaging the RNAi pathway leading to the suppression of target gene expression. RNAi pathway has played a crucial role in our current understanding of genotype to phenotype relations in many organisms, including *C. elegans*. Presence of this pathway in *F. buski* may be useful in deciphering functions of genes as these are not amenable to classical genetic approaches. We tried to decipher from the transcriptome data if this pathway exists in *F. buski* and the divergence from RNAi pathway of other related parasites. The results are summarized in Table 3. Orthologs were found for all the genes associated with small RNA biosynthesis of *C. elegans* in *F. buski* except rde-4. This gene is also absent in the genome of most of the parasitic nematodes except *Brugia malayi* and *Ancylostoma caninum* [65]. The major differences from pathways present in *Schistosoma mansoni* and *C. elegans*, are lack of genes related to dsRNA uptake and spreading (sid-1,sid-2,rsd-6). Absence of these genes is reported in most of the nematodes outside the genus *Caenorhabditis* [65]. In contrast, phylogenetic neighbors of *Schistosoma* sp. possess ortholog of sid-1 which is responsible for dsRNA entry in soaked parasites, though this protein is much larger than sid-1 in *C. elegans*. Another conserved protein rsd-3 is found in almost all the nematodes studied, though its functional collaborators are missing in most of the organisms. [65]. Other than small RNA biosynthesis, proteins associated with nuclear effectors were mostly identified except the absence of less conserved mes-3, rde-2, ekl-5 etc [65]. Overall, orthologs for most conserved proteins from each functional category [65] were found, whereas, the less conserved ones remained unknown.

**Table 3.**
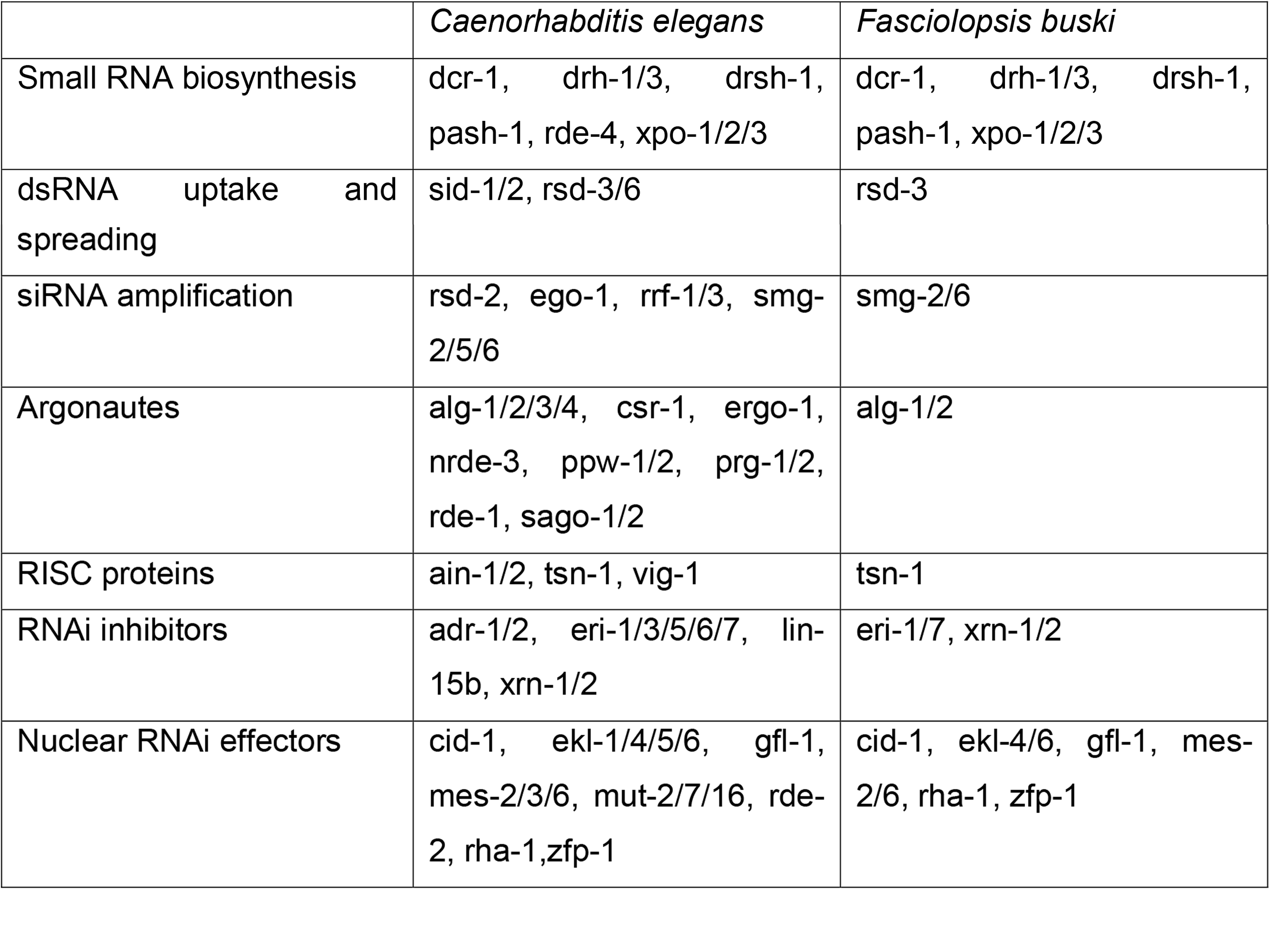
Proteins of RNAi pathway identified from *F. buski transcriptome*.

### Transposable elements (TEs)

Repetitive elements are important structural features of a genome and transcriptome as many of these are transcribed. Retrotransposable elements (RTEs) constitute the main type of interspersed repeats particularly in higher eukaryotic genomes [66]. Therefore, we analyzed the *F. buski* transcriptome for identification of expressed repetitive elements, particularly interspersed elements. A total of 3720 retroelements that include 3477 LINES in the *F. buski* transcriptome are outlined in Table 4. All LINE elements are not expressed; genomic LINE elements were also detected (S9 Table). In contrast to LINES, the data was under-represented for SINE elements. T2#SINE/tRNA were identified from the transcriptome sequences. These results are similar to reports from other trematodes, such as *C. sinensis* [27, 29, 67] where SINE elements were also not reported. Additionally, several LTR retrotransposable elements and DNA transposons (Table 4), whose roles in parasite biology and genome organization are not yet well defined were identified.

**Table 4.**
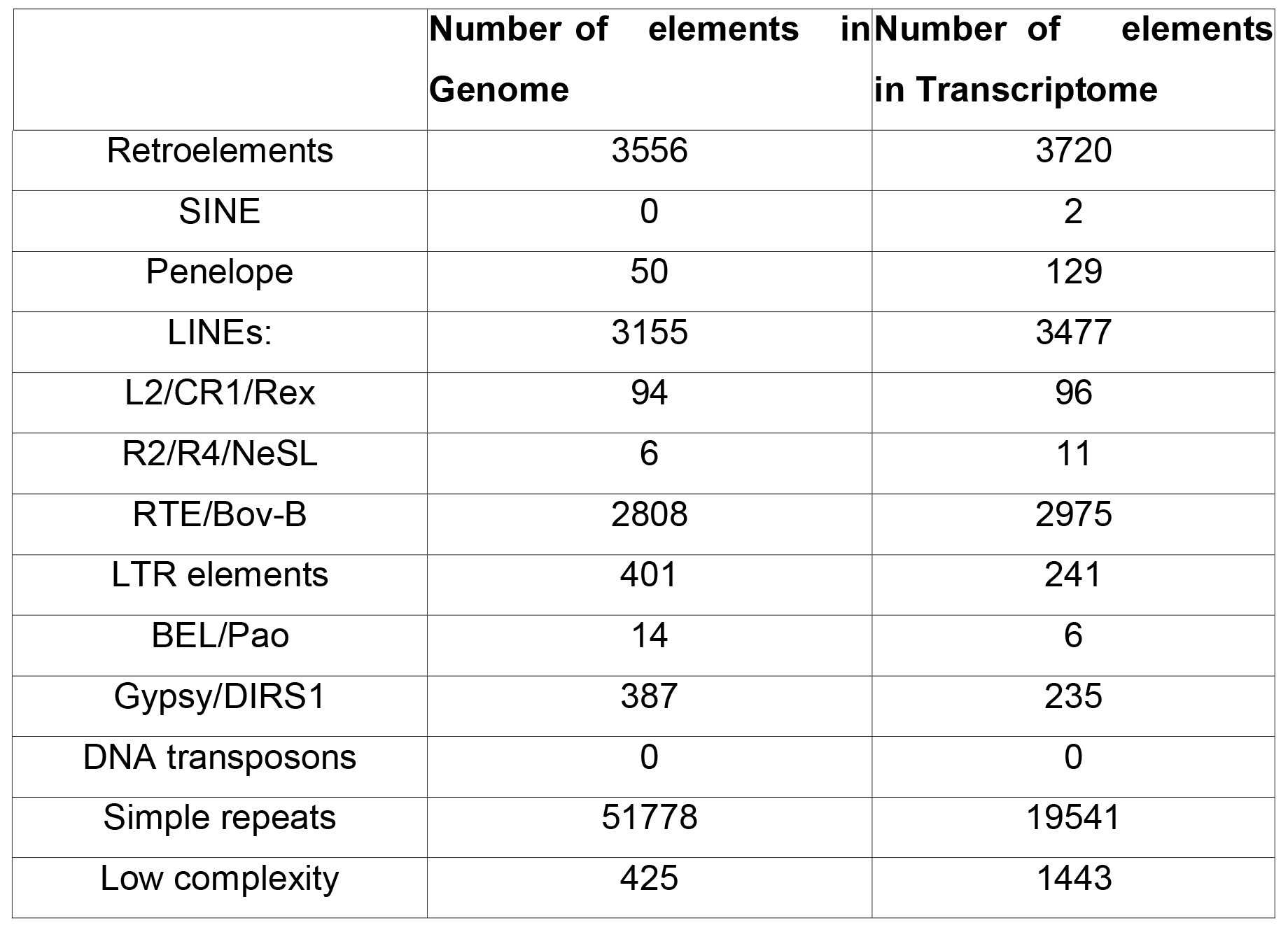
Number of different repeat elements identified from genome and transcriptome of *F. buski*.

|  | Number of elements in<br>Genome | Number of elements<br>in Transcriptome |
| --- | --- | --- |
| Retroelements | 3556 | 3720 |
| SINE | 0 | 2 |
| Penelope | 50 | 129 |
| LINEs: | 3155 | 3477 |
| L2/CR1/Rex | 94 | 96 |
| R2/R4/NeSL | 6 | 11 |
| RTE/Bov-B | 2808 | 2975 |
| LTR elements | 401 | 241 |
| BEL/Pao | 14 | 6 |
| Gypsy/DIRS1 | 387 | 235 |
| DNA transposons | 0 | 0 |
| Simple repeats | 51778 | 19541 |
| Low complexity | 425 | 1443 |

## Discussion

Human parasites are a diverse group of organisms, ranging from unicellular protists to multicellular trematodes. Most well studied molecular data on human host-parasite interactions are from parasites, such as *Plasmodium* and *Leishmania*. Lack of information and complex ecological relationship of multicellular flukes limits our understanding of trematode biology. Recently, availability of new and cheap genome sequencing technologies (NGS) has opened up research avenues for characterization of these organisms. This is reflected in a number of studies that have been published recently on these parasites based on genomic information.

North-east India is considered to be a hot bed of diversity and it is believed that many organisms from this region have evolved interesting biology due to their evolution in a unique ecological niche. In this report we present data about characterization of a trematode parasite *F. buski* (the giant intestinal fluke having a zoonotic potential) using genome sequencing and RNA-seq analysis. Transcriptome sequencing helped us identify 30677 genes from the parasite, and we annotated 12380 genes, driven by longer assembly, that enabled more authentic functional annotation of the unigenes.

Majority of the highly expressed genes in *F. buski* have also been reported in other organisms including trematodes. We note a high level expression of transcripts encoding cytoskeletal elements, protein biosynthesis and folding in the adult stage of the parasite. Parasites infect host and are reliant on the host for their nutrient supply during their growth and development cycle [56]. The parasite is studied at an adult infective stage, where development has switched to the reproductive stage with a high metabolic load required for regular egg production. Reproductive development and egg production require high levels of transcript accumulation and associated protein synthesis, often in conjunction with reorganization of cytoskeletal elements. High-energy costs of eggshell production in *F. buski* require efficient energy generation and high expression of these enzymes may explain this. Cytoskeleton proteins are involved in intracellular transport and glucose uptake in larvae and adult parasites and serve as outlets for their glycogen stores [68]. We noted the high expression of genes encoding fatty acid binding proteins (FABP) that support the role of these molecules in acquisition, storage, and transport of lipids. Since many of these display unique properties, FABPs have been suggested as potential vaccine candidates [69]. We also speculate the difference in the fatty acid metabolic processes, with higher enrichment of gene targets for catabolism, compared to biosynthesis, which can be critical to the parasite biology at the adult stage. Empirical comparison with other data sets from trematodes suggest, absence of biosynthesis in the adult stage, either by a gene-regulatory mechanism or lack of components in the fatty acid biosynthesis. This would warrant further investigation in the light that these parasites have been known to acquire fatty acids from the host [56].

Different proteases are expressed in trematodes such as *F. hepatica*, *C. sinensis* and *S. mansoni*. Proteases play a significant role in parasite physiology as well as in host-parasite interactions [70–72]. In general genes encoding six different ‘catalytic type’ of proteases, were identified in the *F. buski* transcriptome, and were expressed at relatively high-levels. Amongst them, the serine and cysteine proteases and serine protease inhibitors were highly represented in unigenes expressed in the adult stage. In the blood fluke, *S. mansoni*, serine proteases were implicates in tissue invasion and host immune evasion [73, 74]. Cysteine proteases have roles in nutrition, tissue/cell invasion, excystment/encystment, exsheathment and hatching, protein processing and immune-evasion [75]. Papain-like cysteine proteases, particularly the cathepsins, facilitate skin and intestine infections, tissue migration, feeding and suppression of host immune effector cell functions [76]. Serine protease inhibitors (serpins) are a super family of structurally conserved proteins that inhibit serine proteases and are involved in many important endogenous regulatory processes and possible regulation of host immune modulation and/or evasion processes [77].

We report the expression of globin genes (Hemoglobin F2 and Myoglobin I), which was also found to be abundant in *F. buski* transcriptome. The functional role of trematode haemoglobin (Hbs) is still not clear, as adult parasitic trematodes and also nematodes, such as *Ascaris suum*, are residents of semi-anaerobic environment. It is believed that the function of hemoglobin is not only restricted to O_2_ transport but also as an oxygen scavenger, a heme reserve for egg production, and as co-factor of NO deoxygenase [78–82].

The kinome of *F. buski* is similar to other eukaryotes, although the gene family redundancy within protein kinases identified was fewer than reported from free-living nematode. The functional repertoire suggest the presence of an extensive signaling system, that may be required for sensing and modulating signal response in the changing life-cycle and adult development/reproduction in the intra-host environment. CMGC kinases were relatively the most abundant in terms of number of sequences, followed by CAM kinases. This data correlates well with the observation of liver parasite *F. gigantica* [29] and blood fluke, *Schistosoma mansoni* [83]. CMGC kinases are known to regulate cell proliferation and ensure correct replication and segregation of organelles in many eukaryotic organisms including *Plasmodium falciparum* [84]. CAM kinases are associated with calcium-mediated signaling. Among CAMKs in *F. buski*, the majority of kinases belonged to the CAMK-like kinases (CAMK family 1 and 2), death-associated protein kinases (DAPKs) and the myosin light chain kinases (MLCKs). The primary function of MLCK is to stimulate muscle contraction through phosphorylation of myosin II regulatory light chain (RLC), a eukaryotic motor protein that interacts with filamentous actin [83]. The high abundance of the above-mentioned kinase groups may reflect high mobility and muscle activity in *F. buski* associated with feeding and egg sheading.

Expression of genes involved in RNAi pathway suggests partial evidence of an active pathway. We noted absence of the protein required for RNA uptake, and speculate that RNAi would be active if introduced into the cytoplasm considering siRNA as a possible therapy.

*F. buski* genome also encodes different types of transposons as interspersed repetitive sequences similar to other eukaryotes. Members of the Phylum Platyhelminthes are also thought to contain diverse TEs, which comprise up to 40% of their genomes. A total of 29 retrotransposons, belonging to one non-long terminal-repeat (LTR) family (6 elements in CR1) and 3 LTR families (5 elements in Xena and Bel, and 13 elements in Gypsy) have been isolated from the genomes of the digenean trematodes, *C. sinensis* and *Paragonimus westermani*. CsRn1 of *C. sinensis* and PwRn1 of *P. westermani* are novel retrotransposons, which are evenly distributed throughout their genomes and expressed as full transcripts in high copy numbers. Phylogenetic studies have revealed that the CsRn1 and PwRn1 elements formed a novel, tightly-conserved clade, that has evolved uniquely in the metazoan genomes. Diverse retrotransposon families are present in the lower animal taxa, and that some of these elements comprise important intermediate forms marking the course of evolution of the LTR retrotransposons. Retrotransposons in trematodes might influence the remodeling of their host genomes [60].

Two novel families of tRNA-related SINEs have been described to be widespread among all *Salmonoidei* genomes, with a role in human helminth pathogen—*Schistosoma japonicum* (Trematoda: Strigeiformes) [85]. We could identify only two SINE elements from the *F. buski* transcriptome data. Lack of widespread presence of SINE elements in flukes raises interesting questions about evolution of SINEs [85]. SINEs have been reported in early branching protists, such as *Entamoeba histolytica* [86] The inability to detect SINE elements in other trematodes and presence of two families in *F. buski* suggest that these may have come by horizontal transfer possibly with some evolutionary advantage. It is possible that SINEs may have been lost from the genome during evolution in these organisms. We also found that not all LINE elements are expressed in the adult stage. Overall the transcriptome of *F. buski* has shed new light on the biology of this organism.

In conclusion, the rough draft genome and transcriptome characterized in the present study will assist in future efforts to decode the entire genome of *F. buski*. The transcriptome data of the adult stage of this giant intestinal fluke is reported for the first time, and is archived in the NCBI SRA as well as in the database (North-East India Helminth Parasite Information Database) developed by our group (NEIHPID) [87]. We hope our study adds substantially to the public information platform to achieve a fundamental understanding of the parasite biology, which in turn would help in identification of potential drug targets and host-pathogen interaction studies.

## Author Contributions

DKB and VT conceived and designed the experiments, analyzed the data, contributed reagents/materials/analysis tools, wrote the paper. DKB, TR and PP performed the experiments. DKB, TR and PP performed the bioinformatics analysis. All authors reviewed the manuscript.

## Additional Information

The authors declare no competing financial interests.

## DNA Deposition

DNA sequences were deposited as follows:

National Centre for Biotechnology Information (NCBI) Bioproject Accession: PRJNA212796 ID: 212796

NCBI Sequence Read Archive (SRA): **SRP028107**, SRX327895, SRX326786, SRX316736

## Acknowledgement

We would like to acknowledge Dr. Sriram Parameswaran, Genotypic Technology (P) Ltd., Bangalore, Karnataka for his useful suggestions while carrying out the NGS analysis, Dr. Sudip Ghatani, Department of Zoology, NEHU, Shillong for collecting the biosamples and M/s Genotypic Technologies, Bangalore, India for carrying out NGS sequencing for this project.

## Supporting information

**S1 Fig. Histogram displaying length of the unigenes.** Black bars represent all unigenes while red ones represent annotated unigenes only. Clearly, mean of length of annotated unigenes is higher than overall mean length of all unigenes.

**S2 Fig. Major components of metabolic pathways present in F. buski transcriptome.** Colored edges represent proteins homologous to any F.buski unigene.

**S1 Table. Assembly statistics for *F. buski* genome using three denovo assemblers and transcriptome using Trinity.**

**S2 Table. Assembly assessment of uigenes used for annotation**

**S3 Table.** BLASTx hits against NR database

**S4 Table. Top 30 conserved domain/families identified by InterProScan from F. buski transcriptome**

**S5 Table. COG categories assigned to unigenes**

**S6 Table. RPKM values of each of the unigenes**

**S7 Table. List of different kinases identified from *F. buski* transcriptome**

**S8 Table. List of different protease and protease inhibitors identified from *F. buski* transcriptome**

**S9 Table. List of three LINE elements**

